# Covalently Functionalized Click Alginate Cryogels Enable Non-Invasive Near-Infrared Sensing of Gastric pH

**DOI:** 10.64898/2026.09.11.751041

**Authors:** Katia Cherifi, Arturo Israel Machuca Parra, Nahyun Kwon, Lili Ding, Loic Bart, Mouna Ayad, Christopher Rose, Gang Zheng, Simon Matoori

## Abstract

Gastric acidity shapes mucosal injury in reflux disease, peptic ulcer, and gastritis, yet is almost never measured. Acid-suppressive therapy, taken by approximately 40% of elderly adults, is therefore titrated on symptoms alone. A gastric sensor must hold onto its reporter in a highly acidic protease-rich environment, conditions that destabilize ionically crosslinked hydrogels and leach physically entrapped dyes. Click alginates, established in biomechanics and drug delivery but not as sensors, meet this requirement with a single chemistry: bioorthogonal tetrazine- norbornene crosslinks are stable in acid and against proteases, and excess norbornene from crosslinking becomes the anchoring site for the pH-sensitive reporter, so network and reporter are installed by the same mechanism. Cryogelation adds macroporosity for rapid gastric fluid influx. A screen identifies IRDye 680RD as the only near-infrared dye responsive across the targeted pH 2-5. Dye release stays below 1.5% in simulated gastric and intestinal fluids, and the pH response remains active after nine months of storage. In rats, the orally administered NIR fluorescent cryogel resolves pH 2.0 from pH 5.0 transabdominally and discriminates esomeprazole- from saline-treated animals, with extra-gastric organ and plasma fluorescence comparable to untreated controls. *Ex vivo* photoacoustic imaging shows pH-dependent contrast, pointing toward deeper-tissue detection. These results establish covalently functionalized cryogels as a non-invasive sensing platform for hostile physiological environments.

## 1. Introduction

Gastric acidity is a central physiological parameter of the upper gastrointestinal tract, governing food digestion, mineral absorption, antimicrobial defense, and the dissolution and bioavailability of orally administered drugs.^[1]^ Intragastric pH is clinically relevant across a range of acid-related disorders including gastroesophageal reflux disease (GERD), peptic ulcer disease, and *Helicobacter pylori*-associated gastritis where acid secretion shapes symptom burden, mucosal injury, and disease progression.^[1d, 2]^ It also modulates the performance of drugs with pH- dependent solubility and absorption. ^[3]^ Yet despite this importance, gastric pH is rarely measured in routine clinical practice, and therapeutic decisions are typically made empirically rather than from direct acid readouts. ^[1a]^ This gap is particularly consequential for acid- suppressive therapy. Proton pump inhibitors (PPIs) are among the most widely used prescription and over-the-counter drugs, with regular use reported in approximately 15% of the general population and 40% of individuals aged 70 years and older; ^[4]^ and 23.4% of adults worldwide report having used a PPI, ^[5]^ highlighting the risk of overutilization. ^[6]^ However, prolonged use has risks. ^[2e-g, 7]^ By inhibiting the gastric *H^+^/K^+^*-ATPase, PPIs reduce acid secretion and can increase fasted gastric pH from approximately 1.7 to as high as 7.3. ^[8]^ Sustained acid suppression has been linked to nutrient malabsorption, altered drug exposure, and perturbation of the intestinal microbiota.^[2g]^ These risks argue for a more individualized approach in which therapy is titrated to direct measurements of gastric acidity rather than to symptoms or fixed dosing schedules, an approach that would be especially valuable where the balance between acid suppression and physiological gastric function must be carefully maintained.

Existing tools for gastric or esophageal pH assessment are poorly suited to routine, point-of-care use. Catheter-based pH monitoring is invasive, uncomfortable, and disruptive to daily activity, ^[9]^ whereas wireless capsules improve tolerability but remain constrained by cost, workflow demands including capsule recovery, and dependence on specialized equipment. ^[1c, 10]^ More recent ingestible sensors demonstrate the feasibility of monitoring gastrointestinal parameters *in vivo*, yet many are technically complex, require endoscopic or otherwise invasive deployment, or are not optimized for low-cost outpatient use. ^[10–11]^ A simple, broadly deployable platform for repeated pH measurement at the point-of-care is lacking.

Hydrogels are emerging as versatile biosensing materials for biomedical applications.^[12]^ For gastric applications, hydrogels require particular properties to withstand the low pH and high protease activity. ^[13]^ Existing hydrogel-based optical pH sensors typically immobilize the indicator either by physical entrapment, which is prone to dye leaching, or by covalent copolymerization of indicator dyes derivatized with polymerizable groups during network formation. ^[14]^ However, such copolymerization strategies commonly rely on free-radical polymerization, often initiated photochemically or using redox catalysts; the associated radical- generating conditions can produce reactive oxygen species and may compromise biocompatibility or dye stability. Other hydrogel-based gastric monitoring systems rely on embedded electronic components, such as ingestible hydrogel devices carrying wireless sensors, ^[15]^ or surgically implanted hydrogel-circuit sensors that transduce pH through swelling with response times of approximately 1 h. ^[16]^ These electronic systems, however, introduce components that must withstand the corrosive, acidic gastric environment; metallic contacts, batteries, and other electronic elements are vulnerable to corrosion, fouling, and moisture ingress, and battery exposure to gastric fluid raises safety concerns.

Here, we report a minimally releasing, macroporous click-alginate cryogel bearing a covalently immobilized near-infrared (NIR) pH-sensitive dye for fluorescence-based gastric pH sensing (**Figure 1**), an all-polymeric, electronics-free alternative to existing gastric monitoring approaches. Click alginates are hydrogel-forming alginate derivatives with complementary click groups for bioorthogonal covalent crosslinking. ^[17a]^ Unlike these approaches, a single bioorthogonal chemistry here performs both structural and sensing functions: the tetrazine- norbornene reaction that crosslinks the cryogel also anchors a tetrazine-modified commercial NIR dye to residual norbornene groups after gelation, under mild, catalyst-free aqueous conditions, ^[17]^ without dye copolymerization, embedded electronics, or implantation. Because the reporter is introduced after network formation, the same scaffold can, in principle, accommodate any tetrazine-bearing probe. ^[18]^ Using bioorthogonal, highly stable tetrazine- norbornene click chemistry, we anchor the dye within an alginate scaffold to minimize dye release while preserving pH-responsive fluorescence across the relevant gastric pH range. Cryogelation at subzero temperatures generates macroporous networks. ^[19]^ The resulting cryogel shows strong, reproducible pH-dependent fluorescence and minimal dye release, supporting its stability in the gastric environment. Due to their high biocompatibility, hydrophilicity, low immunogenicity, and tunable pore size, click alginate hydrogels are broadly used in biomechanics, tissue engineering, and drug delivery. ^[17a, 20]^ However, their application as biosensing materials is currently underexplored. ^[12]^ Click alginate hydrogels are particularly suited for gastric sensing due to their resistance to low pH and proteases and their click groups allowing for covalent tethering of biosensing molecules. ^[17b, 21]^ When prepared by cryogelation, their macroporosity promises to enable rapid influx of gastric fluids. ^[20a, 20f]^

**Figure 1.**
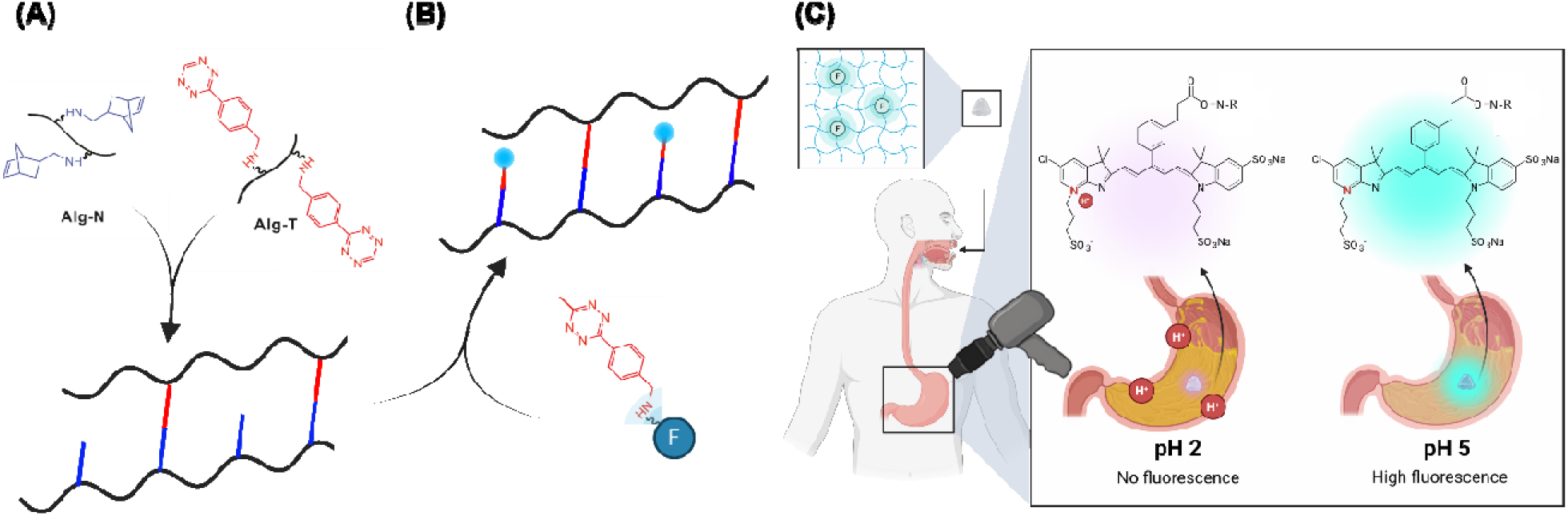
Design of the click alginate cryogel pH sensor for gastric pH monitoring. **(A)** Norbornene-modified alginate (Alg-N) and tetrazine-modified alginate (Alg-T) form a cryogel with a covalently crosslinked macroporous network that allows rapid fluid uptake. **(B)** As Alg-N is used in excess, a tetrazine-functionalized NIR pH-sensitive dye (IRDye 680RD) is coupled to residual norbornene groups on the network, resulting in covalent anchoring of the probe in the cryogel. **(C)** In clinical use, the pH-sensitive cryogel is administered orally and changes its optical properties in relation to gastric fluid pH which could be detected by a non-invasive portable detector.

## 2. Materials and Methods

### 2.1 Materials

High molecular weight alginate (I-1G, 280 kDa ^[22]^) was obtained from Kimica Corporation (Tokyo, Japan). (4-(1,2,4,5-Tetrazin-3-yl)phenyl)methanamine hydrochloride salt (benzylamino tetrazine) was purchased from KareBay™ Biochem, Inc. (Buffalo, NY) for click-functionalized alginate polymer synthesis, and (4-(6-methyl-1,2,4,5-tetrazin-3-yl)phenyl)methanamine hydrochloride salt (methyltetrazine amine) was obtained from BroadPharm (San Diego, CA) for dye-linker conjugation. 5-Norbornene-2-methanamine, purified pepsin, sodium hydroxide, and Spectrum™ Labs Spectra/Por™ 12-14 kDa MWCO dialysis kits were purchased from Fisher Scientific (Pittsburgh, PA). Activated charcoal, sodium chloride, hydrochloric acid, potassium phosphate monohydrate, dimethyl sulfoxide, 1-ethyl-3-(3-dimethylaminopropyl) carbodiimide hydrochloride, *N*-hydroxysuccinimide, 2-(*N*-morpholino) ethanesulfonic acid, trisodium citrate dihydrate, 8-hydroxypyrene-1,3,6-trisulfonic acid (HPTS), 5(6)-carboxyfluorescein, pancreatin from porcine pancreas (4× USP specifications, ≥100 USP Units of protease activity per mg; catalogue no. P1750), and Nalgene® vacuum filtration system (0.22 µm) were purchased from Sigma-Aldrich (St. Louis, MO). Citric acid was purchased from A&C American Chemicals Ltd. (Saint-Laurent, QC, Canada). IRDye 680RD NHS ester (IRDye 680RD) and IRDye 800CW NHS ester (IRDye 800CW) were purchased from LI-COR Biotech, LLC (Lincoln, NE, USA). Sulfo-cyanine 7 carboxylic acid (sulfo-cyanine 7), sulfo-cyanine 7.5 carboxylic acid (sulfo- cyanine 7.5), indocyanine green (ICG), AF 647 NHS ester (AF647), and BDP® 630/650-X-NHS ester (BDP) were purchased from Lumiprobe (Westminster, MD). Esomeprazole sodium (Nexium® i.v., 40 mg lyophilized powder for solution for injection/infusion) was obtained from Grünenthal Pharma AG (Glarus Süd, Switzerland). Teklad™ 2916 low-fluorescence rodent diet was purchased from Envigo (Indianapolis, IN).

### 2.2. Methods

#### 2.2.1 Fluorescent dye screen for pH sensitivity

To evaluate the pH sensitivity of the free dyes, each fluorophore was dissolved in 12.5 mM trisodium citrate buffer and adjusted to pH values from 1.0 to 6.0. The dye panel consisted of 5(6)-carboxyfluorescein, HPTS, AF647, IRDye 800CW, IRDye 680RD, sulfo-cyanine 7, sulfo-cyanine 7.5, ICG, and BDP. Dye concentrations were chosen to yield absorbance signals between approx. 0.1 and 1.5: 5 µM for sulfo-cyanine 7; 10 µM for AF647, IRDye 800CW, IRDye 680RD, and sulfo-cyanine 7.5; 25 µM for ICG and BDP; 0.2 mM for HPTS; and 50 µM for 5(6)- carboxyfluorescein. For all dyes, 150 µL of solution was added to clear, flat-bottom 96- well microplates (Greiner; Sigma-Aldrich, St. Louis, MO), and absorbance spectra from 400 to 900 nm were recorded on a Spark multimode microplate reader (Tecan, Männedorf, Switzerland).

#### 2.2.2 Preparation of dye-conjugated click alginate cryogels

##### Click functionalization of alginate hydrogels

Click-functionalized alginates were synthesized by covalent modification with either 5- norbornene-2-methanamine or benzylamino-tetrazine as previously described. ^[17a]^ High- molecular-weight sodium alginate was dissolved to 0.5% (w/v) in buffer containing 0.1 M 2-(N- morpholino) ethanesulfonic acid (MES) and 0.3 M NaCl at pH 6.5. 2.2 grams of N- hydroxysuccinimide (NHS) and 3.7 grams of 1-ethyl-3-(3-dimethylaminopropyl) carbodiimide hydrochloride (EDC) per gram of alginate were added, followed by either norbornene or tetrazine (1 mmol/g alginate) to give norbornene-modified (Alg-N) or tetrazine-modified (Alg-T) derivatives, respectively. Coupling proceeded under continuous stirring at room temperature for 24 h. The reaction mixture was centrifuged at 4700 RPM for 15 min at room temperature and supernatants were combined and dialyzed in 12-14 kDa MWCO tubing against distilled water using a stepwise decreasing NaCl gradient from 150 mM to 0 mM, with intermediate steps at 0.1 M and 0.05 M, the dialysis solution changed every 2 h, and each salt concentration maintained for 4 h before progressing to the next step. Dialyzed solutions were treated with activated charcoal, sterile filtered (0.22 µm), lyophilized, and stored at -20 °C.

Alg-T was dissolved in PBS (50 mM, isotonic, pH 7.4) at 10 mg/mL and Alg-N at 15 mg/mL; both solutions were vortexed. Equal volumes (30 µL) of Alg-T and Alg-N were rapidly combined in a 1.5 mL microcentrifuge tube to initiate click crosslinking, immediately vortexed, and centrifuged. Precursor mixtures were transferred to -20 °C immediately and incubated for 24 h for cryogelation (**Figure 2**).

**Figure 2.**
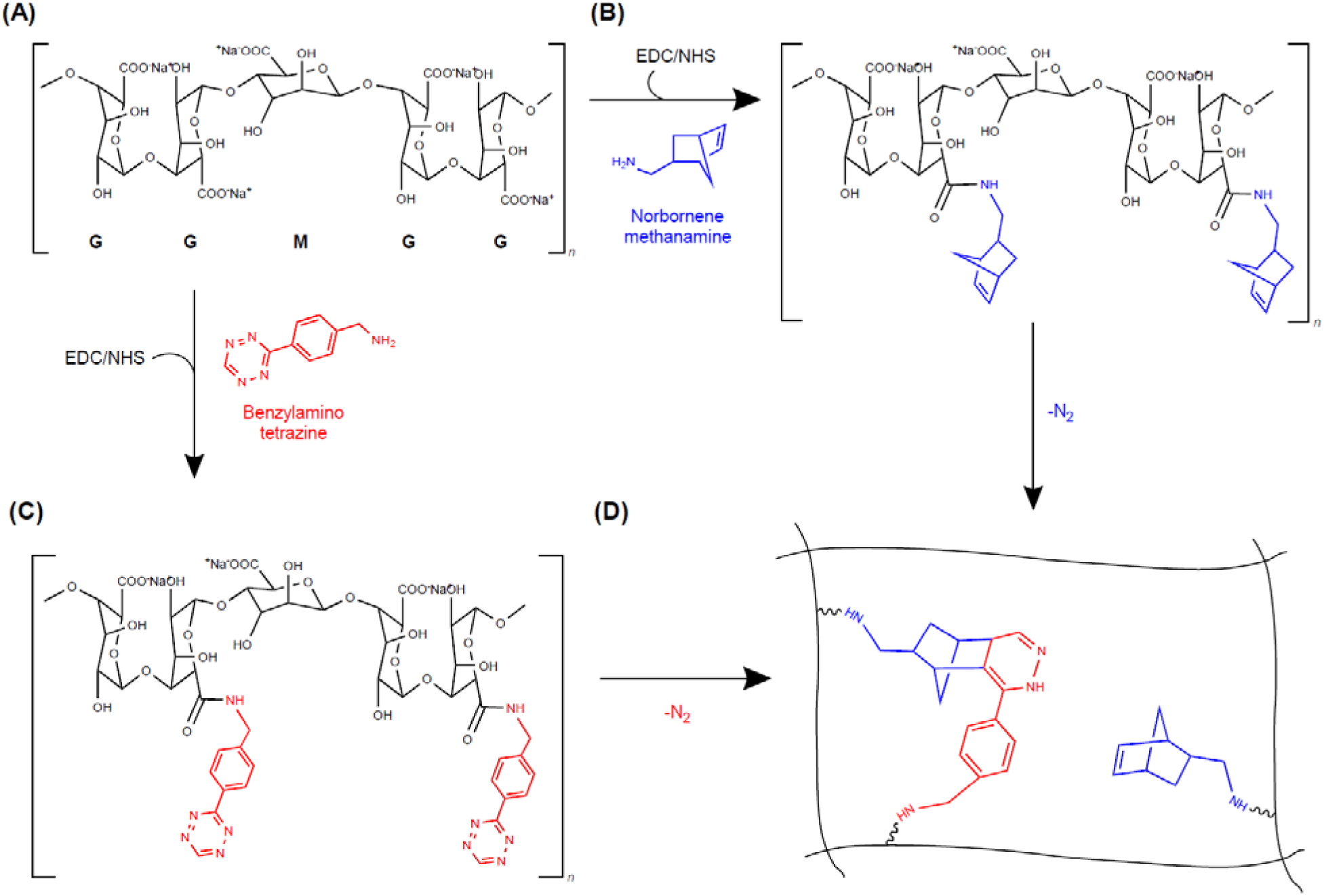
Click functionalization of alginate polymer. **(A)** Native alginate (M/G ≈ 0.5, ∼60-70% G content); **(B)** Norbornene-modified alginate; **(C)** Tetrazine-modified alginate; **(D)** Crosslinked cryogel formed by inverse electron-demand Diels-Alder (IEDDA) cycloaddition between (B) and (C). For clarity, functional groups are drawn on G-unit carboxyls, but modification occurs at carboxyls along both M and G segments (approximately 5% degree of substitution, as previously reported for this synthesis protocol). ^[17a]^

##### Synthesis of dye-linker conjugate

Methyltetrazine-amine hydrochloride was dissolved in 50 mM isotonic PBS at pH 7.4 at 1 mg/mL and diluted in PBS to stock solutions of 90 µM or 450 µM. IRDye 680RD NHS ester was diluted from a stock (0.9 mM) in DMSO to 90 µM or 450 µM in 50 mM isotonic PBS at pH 7.4. Equal volumes of the dye and methyltetrazine-amine stock solutions of the same concentration were combined on ice (1:1 molar ratio; 20 µL total), yielding final concentrations of 45 µM or 225 µM of each component, and incubated at 4 °C on an orbital shaker for 24 h, protected from light, to allow amide bond formation. To determine the dye loading required for cryogel functionalization, conjugates prepared from 90 µM and 450 µM stock solutions were initially compared against unfunctionalized cryogels (**Supporting Figure 1**); all subsequent experiments were performed using the conjugate prepared from 90 µM stock solutions.

##### Conjugation of dye-linker to click-hydrogels

Frozen Alg-T/Alg-N cryogels were thawed at room temperature for 2 h prior to functionalization. The dye-tetrazine conjugate prepared (20 µL, 45 µM) was added to individual cryogels to initiate tetrazine-norbornene ligation, giving a total functionalization volume of approximately 80 µL and a final concentration of 11.25 µM IRDye 680RD and methyltetrazine-amine in the cryogel (**Figure 3**). Samples were incubated at 37 °C on an orbital shaker for 24 h, protected from light.

**Figure 3.**
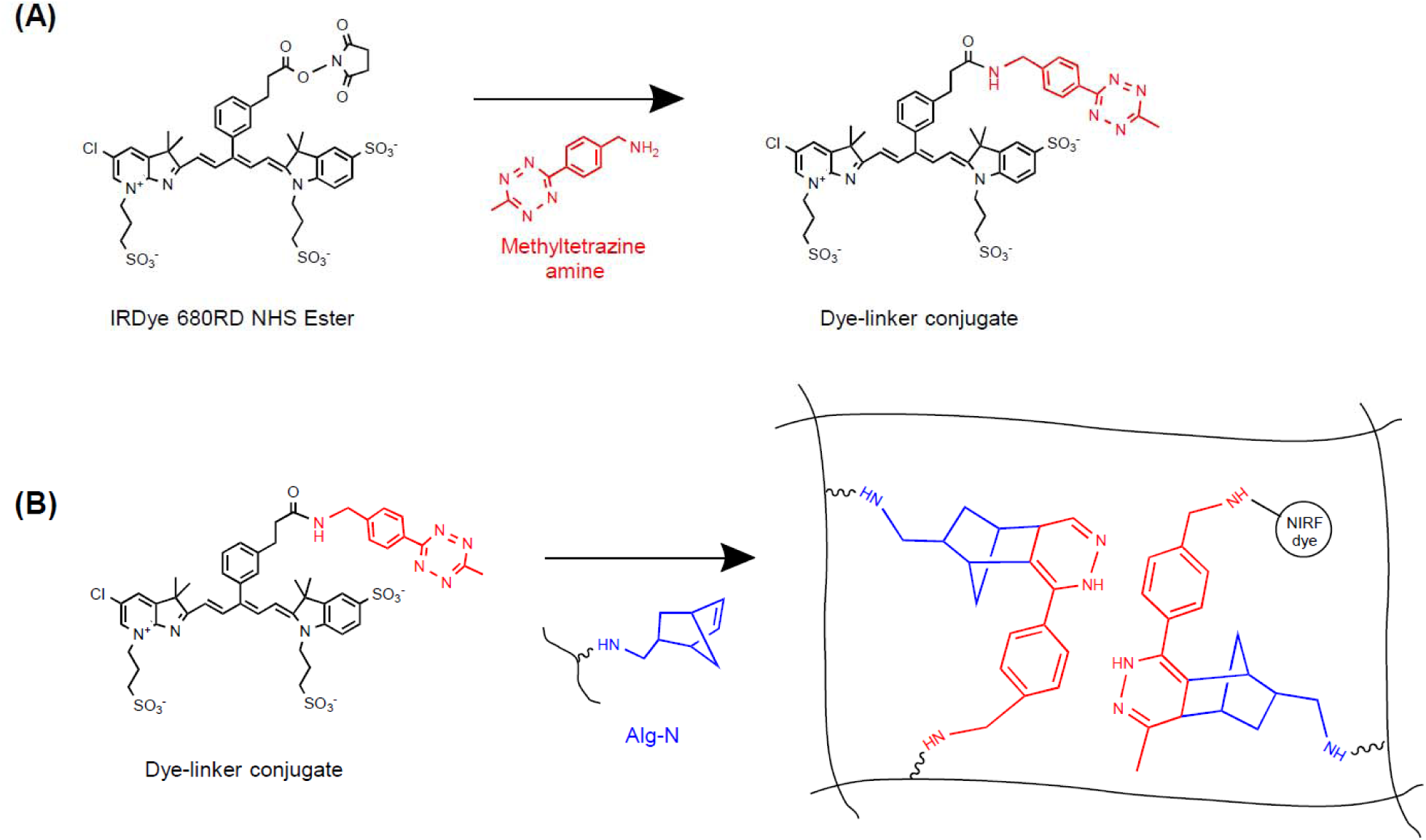
Click functionalization of the NIR dye and conjugation to the click alginate cryogel. **(A)** IRDye 680RD NHS ester reacts with methyltetrazine-amine to form a dye-tetrazine conjugate; **(B)** the conjugate is covalently attached to the norbornene-modified network by IEDDA cycloaddition between the dye’s tetrazine and excess norbornene groups.

Functionalized cryogels were purified by dialysis. Each cryogel was immersed individually in 40 mL of deionized ultrapure water and agitated at 4 °C for 3 d, with daily replacement of the dialysis medium to remove unbound conjugate and reaction by-products. Purification progress was monitored by fluorescence. Cryogels were then transferred to black 96-well plates and fluorescence recorded at an excitation wavelength of 664 nm and emission wavelength of 702 nm (20 nm bandwidths each). An 11 × 11 read grid (121 reads per well) was performed for each well, with blank well controls subtracted to isolate hydrogel-specific signal values.

#### 2.2.3. Cryogel characterization

##### Macroporous structure of click alginate cryogel

Macroporous structure of the dye-conjugated click alginate cryogel in Milli-Q water was evaluated by fluorescence microscopy using a ZEISS Axio Observer microscope equipped with an X-Cite 120 LED mini-illumination system, an Axiocam MRm camera, and a ZEISS Filter Set 50 (excitation: BP 640/30 nm; dichroic beam splitter: FT 660 nm; emission: BP 690/50 nm). Cryogels were cut into thin sections to enable transmission of the excitation light and visualization of the porous structure. Image analysis was conducted using ZEN Microscopy Software (ZEISS).

##### Mechanical properties of click alginate cryogel

The mechanical properties of the dye-conjugated click alginate cryogels were evaluated by unconfined compression using a TA. XT Plus Texture Analyzer (Stable Micro Systems, Surrey, UK). ^[23]^ Cryogels were placed on an aluminum platform and compressed vertically using a 10 mm-diameter cylindrical stainless-steel probe at a speed of 0.0167 mm/s (1 mm/min) to a maximum penetration depth of 1.5 mm. Initial cryogel heights were determined from the axial gap of the rheometer prior to compression testing. Force-distance profiles were recorded for three independently prepared cryogels. Compressive stress was calculated by dividing the maximum force at 1.5 mm displacement by the area of the cryogel’s circular face (5 mm diameter).

Rheological properties of the hydrogel sensor were evaluated using a Discovery HR-20 rheometer (TA Instruments, New Castle, DE) following established oscillatory rheology protocols for hydrogels. ^[24]^ Cryogels were loaded between 8 mm parallel plates at a gap adjusted to achieve an axial force of approximately 0.01 N, and frequency sweeps were performed from 0.1 to 10 rad/s (ω ≈ 0.016-1.6 Hz) at a constant strain of 1% at room temperature, a strain within the linear viscoelastic region of the cryogels as determined in preliminary amplitude sweeps. ^[24–25]^

#### 2.2.4 *In vitro* evaluation of dye-conjugated click alginate cryogel activity and stability

##### Activity and stability in simulated gastrointestinal fluids

The pH-dependent fluorescence of the dye-conjugated cryogel was evaluated against that of the dye-tetrazine conjugate in trisodium citrate buffer and in simulated gastric fluid (SGF). The dye- tetrazine conjugate was measured in 12.5 mM trisodium citrate buffer across pH 1.0-6.0 to establish a baseline response. Purified dye-conjugated cryogels were placed in black 96-well plates, with unfunctionalized cryogels used as blanks for background subtraction. Cryogels were equilibrated in 12.5 mM trisodium citrate buffer (pH 1.0-6.0) at room temperature, protected from light, for 15 min before measurement. SGF was prepared by adding purified pepsin (3.2 mg/mL) to 12.5 mM trisodium citrate buffer at pH 1.0-6.0 in accordance with the literature. ^[26]^ The pH response was evaluated for the dye-tetrazine conjugate and for the dye-conjugated cryogel as above. Cryogel stability and dye release were also quantified in simulated intestinal fluid (SIF), prepared from 50 mM potassium phosphate buffer at pH 6.8 containing porcine pancreatin (17.5 mg/L; Sigma-Aldrich, P1750, 4× USP specifications), corresponding to 1750 U/L protease activity. ^[26d, 26e, 27]^ Dye-conjugated cryogels were incubated at 37 °C in 1 mL of SGF (pH 2.0) or SIF; individual cryogels (n = 3) were incubated on a rotating platform. At 0.5, 1, 2, 4, and 24 h, a 100 µL aliquot of the supernatant was sampled for fluorescence measurement, and the cryogel was transferred to a fresh tube containing 1 mL of the corresponding fresh medium. Released amounts were quantified against calibration curves for the dye-tetrazine conjugate in SIF or SGF, corrected for the sampled volume and any dilution. Cumulative release was calculated as the sum of dye released across all intervals and expressed as a percentage of the total dye content retained in each cryogel after washing. Long-term stability was assessed by measuring fluorescence at pH 2.0 and pH 5.0 in 300 mM trisodium citrate buffer, as described below. Two batches prepared 9 months apart and stored at 4 °C in Milli-Q water, protected from light, were tested in parallel, with fluorescence measured at pH 2.0 and pH 5.0 in 300 mM trisodium citrate buffer.

#### 2.2.5 *In vivo* evaluation of dye-conjugated click alginate cryogel

All procedures were approved by the institutional animal ethics committee at the CrCHUM (Montréal, QC, Canada) under protocol numbers CM20020CRr (pH 2.0 versus pH 5.0 study) and CM24016CRr (esomeprazole versus saline study) and were conducted in accordance with the guidelines of the Canadian Council on Animal Care (CCAC). Male Sprague-Dawley rats (CD® (SD) IGS) of 8-12 weeks of age obtained from Charles River, used in the *in vivo* studies.

##### Activity testing in healthy rats gavaged with buffers at pH 2.0 or 5.0

We first evaluated the dye-conjugated click alginate cryogel’s fluorescence response in healthy rats receiving 300 mM trisodium citrate buffered solutions at pH 2.0 or pH 5.0. The rats were fasted for 14 h with free access to water. The abdomen was shaved and each animal weighed to confirm compatibility with the imaging platform (≤ ∼500 g). During the experiment, rat feces were removed to avoid coprophagy. Rats were anesthetized with isoflurane (3% induction, 2% maintenance in oxygen) for 15 min, and a baseline (control) scan of the gastrointestinal tract was acquired on an eXplore Optix MX2 *in vivo* imaging system (Advanced Research Technologies, Saint-Laurent, QC, Canada) with 666 nm excitation and a 693 nm long-pass emission filter. Rats were allowed to recover in the sternal position until they regained normal consciousness. Each rat then received an oral gavage of pure water alone (2 mL) to flush the stomach, was re-anesthetized (3% induction, 2% maintenance) for 15 min, and imaged again under identical settings. After recovery, each rat received an oral gavage of citrate buffer (trisodium citrate 300 mM; pH 2.0 or pH 5.0) containing four finely chopped cryogels in 2 mL total volume per rat at room temperature. Five minutes after administration, animals were re- anesthetized (3%/2%) for 15 min and the abdominal area was imaged as before. Animals were monitored for 12-24 h for respiratory distress or abnormal behavior. Rats were studied using a within-subject crossover design, with each animal receiving both the pH 2.0 and pH 5.0 cryogel- containing buffers in separate sessions. The buffer pH was alternated between sessions for each animal, ensuring that no rat received the same pH condition in consecutive sessions. Body weight and clinical condition were recorded weekly (**Supporting Figure 2**). The mean fluorescence intensity of each gastric region of interest (ROI) was quantified using the imaging software. The fluorescence intensity of rats receiving pH 5.0 was normalized to the mean of all rats receiving pH 2.0. Rats were imaged weekly for at least 4 weeks and until weight increase limited compatibility with the imaging platform (≤ ∼500 g). At study end, animals were euthanized by inhalation of 4% isoflurane until cardiac/respiratory arrest, followed by cervical dislocation, per approved protocols, for post-mortem analysis. Euthanasia and necropsy were performed immediately after the final imaging session, 2 h after the final oral administration.

##### Activity testing in esomeprazole- versus saline-treated rats

We next tested whether the dye-conjugated click alginate cryogel could distinguish normal from pharmacologically elevated gastric pH under chronic PPI therapy. The gel was evaluated in rats receiving either once-daily subcutaneous esomeprazole (20 mg/kg; 20 mg/mL in saline; 1 mL/kg) on weekdays (Monday-Friday) or sterile saline (1 mL/kg) on the same schedule. Treatment and imaging followed a weekly cycle from the start of the study, with imaging performed on Fridays after the injection.

Before imaging, animals were fasted for 14 h with free access to water and the abdomen was shaved. Each rat received an oral gavage of pure water (2 mL) to rinse the stomach, was anesthetized (3%/2%) for 15 min, and imaged on the Optix MX2 system as above. Subsequently, each rat received an oral gavage of four finely chopped cryogels suspended in 2 mL of water and was re-anesthetized (3%/2%) 5 min later for 15 min and imaged under identical parameters. Animals were monitored for 12-24 h after each imaging session, and body weight and health status (respiration, movement/activity, and general clinical condition) were assessed weekly. Fluorescence intensity was quantified as above with normalization of esomeprazole- treated rats to saline controls. Rats were imaged weekly for at least 4 weeks and until weight increase limited compatibility with the imaging platform (≤ ∼500 g). At study end, animals were euthanized by inhalation of 4% isoflurane until cardiac/respiratory arrest, followed by cervical dislocation, per approved protocols, for post-mortem analysis. Euthanasia and necropsy were performed immediately after the final imaging session, 2 h after the final oral administration.

##### *Ex vivo* organ and plasma fluorescence analysis

An untreated healthy, age-matched male Sprague-Dawley rat obtained from the same animal facility, which had not received any treatment, cryogel, or gavage, was included to determine background organ and plasma fluorescence. Because cryogel-treated rats received repeated oral administrations throughout the weekly imaging schedule, organ-associated fluorescence at necropsy reflects cumulative exposure rather than the biodistribution of a single dose. Residual gastric fluorescence may also reflect incomplete clearance of gastric contents during the 2 h interval between the final gavage and necropsy. Rats were euthanized as described above. Blood was collected from the heart under anesthesia prior to cervical dislocation and anticoagulated with heparin. The stomach, spleen, lungs, heart, brain, bladder, liver, and large and small intestines and kidneys were excised. Organs were imaged on the Optix MX2 system using the same acquisition settings as for *in vivo* imaging. Fluorescence intensity was quantified from organ regions of interest and expressed as relative fluorescence intensity for comparison between the untreated control animal and cryogel-treated animals (**Supporting Figure 3**).

Stomachs were formalin-fixed, paraffin-embedded, sectioned, and stained with hematoxylin and eosin (H&E) by the CrCHUM histology platform. Histological evaluation included one control rat, three saline-treated rats, and three esomeprazole-treated rats. Slides were anonymized and evaluated in a blinded manner by a pathologist, who prepared a semi-quantitative histopathology report. Gastric mucosa was scored on a 0-3 scale for epithelial integrity, parietal- cell alteration, inflammation, and necrosis, where 0 indicated no abnormality and 3 indicated severe alteration. Epithelial integrity was assessed based on preservation of the gastric epithelium and the presence and extent of superficial erosions.

Plasma was collected at the terminal time point from untreated control, esomeprazole-treated, and saline-treated rats from blood obtained by cardiac puncture under anesthesia prior to cervical dislocation, as described above. Plasma fluorescence was measured using a Spark multimode microplate reader (Tecan, Männedorf, Switzerland) under the same acquisition settings for all samples and expressed as relative fluorescence intensity for descriptive comparison among groups.

#### 2.2.6 *Ex vivo* photoacoustic imaging study

We next evaluated the dye-conjugated click alginate cryogel performance by photoacoustic (PA) imaging in an *ex vivo* mouse stomach model to explore an alternative modality with greater tissue penetration depth. All mouse procedures were approved by University Health Network animal care committee under AUP 5922. Female BALB/c mice at the age of 8-12 weeks obtained from Envigo (Indianapolis, IN) were used for this study.

Dye-conjugated click alginate cryogels were prepared as described in Section 2.2.2, except that 800 µM stock solutions of IRDye 680RD and methyltetrazine-amine were used instead of 90 µM stocks, increasing the final concentration of both components in the cryogel from 11.25 µM to 100 µM; Alg-T and Alg-N concentrations and volumes were unchanged. Before imaging, each cryogel was incubated in 300 mM trisodium citrate buffer at either pH 2.0 or pH 5.0 for 30 min at room temperature to allow equilibration and to visually confirm the expected pH-dependent color change (**Supporting Figure 4**). Each mouse received 250-350 mg of hydrogel, finely chopped and suspended in 100 µL of the corresponding pH 2.0 or pH 5.0 buffer, yielding a total suspension volume of approximately 350-450 µL. The suspension was loaded into a 1 mL syringe for gastric administration. Five mice received hydrogel suspensions equilibrated at pH 2.0 and five received hydrogels equilibrated at pH 5.0. In addition, one mouse given only pH 2.0 buffer and one mouse given only pH 5.0 buffer were imaged as negative controls to quantify background PA signal in the absence of hydrogel.

Cryogel suspensions were administered to live mice by oral gavage. Mice were then euthanized by 5% isoflurane inhalation until cardiac/respiratory arrest followed by cervical dislocation, and stomachs were excised for immediate *ex vivo* imaging in a PA-ultrasound system. Photoacoustic imaging was performed on a LAZR system (VisualSonic Vevo) and data was acquired at 680 nm with a 21 MHz transducer, PA gain of 43 dB, and B-mode gain of 18 dB, with all acquisition parameters held constant across mice and treatment groups. Gastric regions of interest (ROIs) were defined on the PA images, and the mean PA signal intensity was quantified from these ROIs for each sample. The mean PA signal of the buffer-only negative control at the corresponding pH was subtracted from each sample value to remove background signal in the absence of hydrogel; the resulting background-corrected values are reported.

#### 2.2.7 Statistical analysis

Statistical analyses were performed using GraphPad Prism (version 8.0.1). Fluorescence intensities were normalized as described for each experiment (to the pH 2.0 group or to saline controls for *in viv*o data, and to unfunctionalized cryogels or buffer-only controls for background subtraction). Differences among three or more groups were assessed using one-way or two-way analysis of variance (ANOVA), as appropriate, followed by Tukey’s multiple-comparisons test. Comparisons between two independent groups were performed using two-tailed unpaired *t*- tests. Comparisons within the same animals (crossover design) were performed using two-tailed paired t-tests. Data are presented as mean ± SD. A value of *p* < 0.05 was considered statistically significant.

## 3. Results and Discussion

### 3.1. Selection of a pH-sensitive NIR dye for gastric sensing

To identify a dye responsive in the target gastric pH range, we screened a panel of commercial far-red and NIR fluorophores (**Figure 4**). HPTS and 5(6)-carboxyfluorescein were included as positive controls with known pH dependent optical properties, and showed the expected pH dependence (**Figure 4A, 4B**). ^[23a, 28]^ Despite their pH-dependent absorbance behavior, both reference dyes emit in the visible range, where hemoglobin and other tissue chromophores absorb strongly, limiting their suitability for deep-tissue *in vivo* imaging. ^[29]^

**Figure 4.**
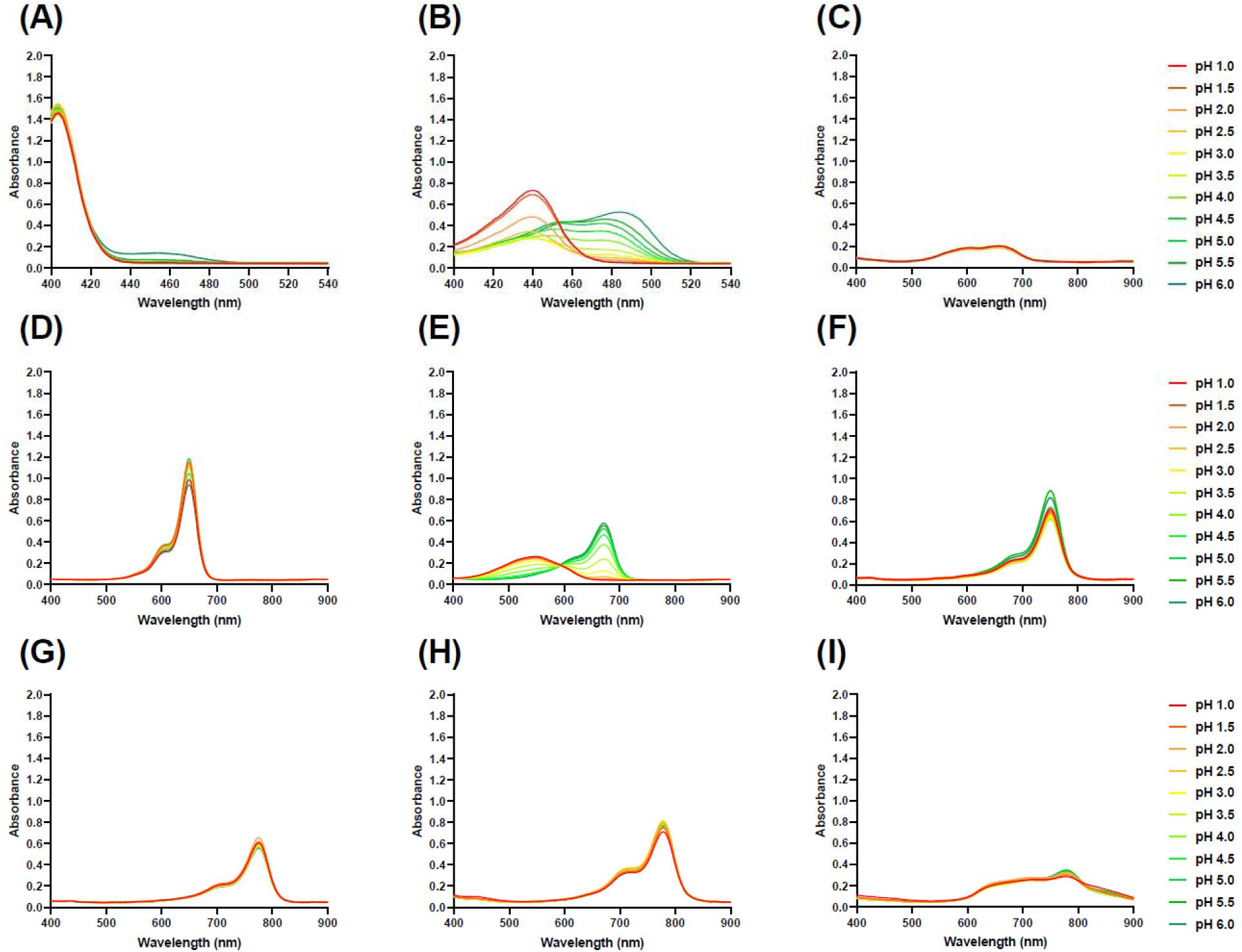
Screen of fluorescence dyes for pH dependence. Absorbance spectra of **(A)** 8- hydroxypyrene-1,3,6-trisulfonic acid (HPTS; 0.2 mM), **(B)** 5(6)-carboxyfluorescein (50 µM), **(C)** BDP® 630/650-X NHS ester (25 µM), **(D)** AF647 NHS ester (10 µM), **(E)** IRDye 680RD NHS ester (10 µM), **(F)** sulfo-cyanine 7 carboxylic acid (5 µM), **(G)** IRDye 800CW NHS ester (10 µM), **(H)** sulfo-cyanine 7.5 carboxylic acid (10 µM), and **(I)** indocyanine green (ICG; 25 µM), measured over pH 1.0-6.0 in 12.5 mM trisodium citrate buffer (*n* = 1).

To identify a fluorescence dye suited for *in vivo* imaging, we screened the following far-red and NIR dyes: BDP, AF647, IRDye 680RD, sulfo-cyanine 7, IRDye 800CW, sulfo-cyanine 7.5, and ICG (**Figure 4C-I**). The far-red dyes BDP 630/650-X and AF647 displayed well-defined far-red absorbance bands but low spectral variation between pH 1.0 and 6.0 (**Figure 4C, D**). Among the NIR dyes, IRDye 680RD was the only dye to show pronounced pH-dependent changes in its NIR absorbance band around 680 nm, including increased peak intensity and spectral shifts from pH 1.0 to 6.0 (**Figure 4E**). Sulfo-cyanine 7, IRDye 800CW, sulfo-cyanine 7.5, and ICG showed largely pH-independent spectra over the same range (**Figure 4F-I**).

The limited pH response of BDP 630/650-X and the cyanine dyes other than IRDye 680RD is consistent with the relative insensitivity of BODIPY and canonical cyanine chromophores to protonation in the examined acidic range, where such protonation does not substantially perturb the conjugated electronic system. ^[30]^ In contrast, IRDye 680RD exhibited pronounced pH- dependent changes in NIR absorbance throughout the clinically relevant gastric pH range of 2.0-5.0 as reported by us in the past, ^[31]^ identifying it as the most promising NIR candidate for gastric pH sensing. pH-dependent protonation of the imine nitrogen on the five-membered ring of the azaindolenine likely modulates π-electron delocalization within IRDye 680RD, producing an increased absorbance with deprotonation at increasing pH. ^[31]^ The reported excitation and emission maxima of approximately 680 and 694 nm, respectively, also make IRDye 680RD well suited for NIR fluorescence.

### 3.2. Design and characterization of the pH-sensing dye-conjugated click alginate cryogel

To translate the identified pH-sensitive NIR dye into a gastric biosensing system with minimal absorption, we first engineered an alginate scaffold functionalized for covalent bioorthogonal conjugation. Building on click alginate chemistry, ^[17a]^ we synthesized alginate-tetrazine and alginate-norbornene, and prepared a macroporous hydrogel by cryogelation. This cryogel bears bioorthogonal handles that can covalently bind ligands carrying a complementary click group. ^[12, 17a, 32]^ Here, we synthesized an IRDye 680RD derivative bearing a tetrazine moiety for covalent conjugation to the click cryogel. The covalent conjugation with an acid-stable bond aims at minimizing dye release to preserve pH responsiveness and minimizing dye absorption. The dye was first functionalized with a tetrazine linker giving a click-compatible derivative that retained pH-responsive NIR fluorescence (**Figure 5A**) while providing a handle for conjugation to the cryogel. Incubating the click alginate cryogel with this dye-tetrazine conjugate led to IEDDA cycloaddition between the dye’s tetrazine and the excess norbornene sites on the network (**Figure 3**), yielding cryogels with significantly higher fluorescence than unfunctionalized controls (**Figure 5B**, **Supporting Figure 1**). Increasing the dye stock concentration fivefold, from 90 µM to 450 µM, resulted in only a modest further increase in cryogel fluorescence (**Supportin Figure 1**). The 90 µM condition was therefore considered sufficient and was used for all subsequent experiments. The dye is thus covalently anchored rather than physically entrapped, minimizing uncontrolled release and systemic absorption and supporting a minimally releasing gastric sensor. This confirms that bioorthogonal chemistry developed for peptide-functionalized alginate can be repurposed for small-molecule NIR probes, broadening the platform toward a wider class of covalently immobilized optical sensors. ^[18, 33]^

**Figure 5.**
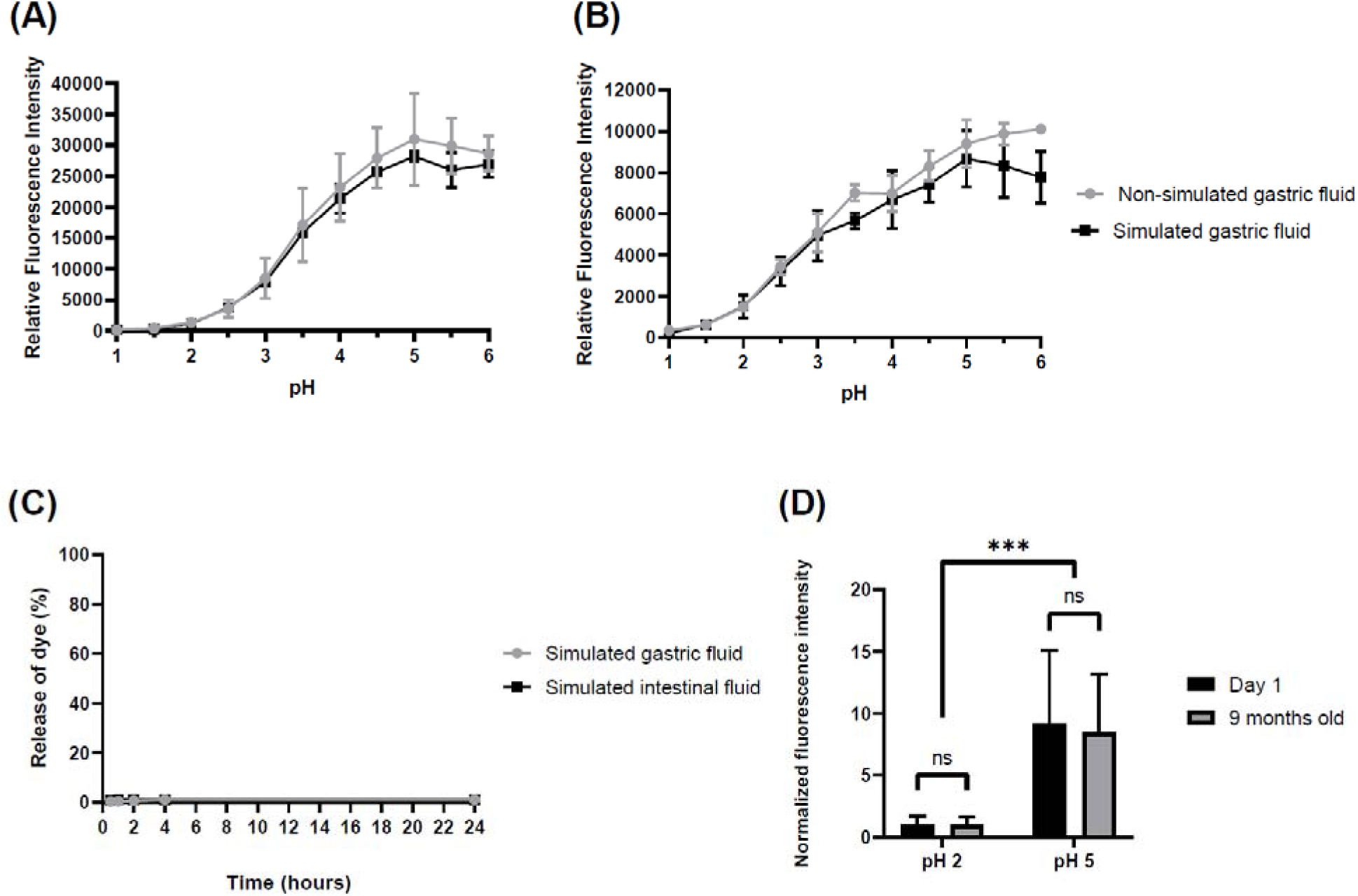
*In vitro* testing of pH-sensitive IRDye 680RD-conjugated click alginate cryogel activity and stability. **(A)** Relative fluorescence intensity of the tetrazine-IRDye 680RD conjugate measured in trisodium citrate buffer and SGF across pH 1.0-6.0. **(B)** Relative fluorescence intensity of dye-conjugated click alginate cryogel measured in trisodium citrate buffer and SGF across pH 1.0-6.0. **(C)** Cumulative release of IRDye 680RD from dye-conjugated click-alginate cryogels incubated in SGF and SIF over 24 h. **(D)** Normalized fluorescence intensity of dye-conjugated cryogels at pH 2.0 and pH 5.0 after storage for one day and 9 months. Data are presented as mean ± SD (*n* = 3). Two-way ANOVA with Tukey’s multiple-comparisons test; pH 2 vs pH 5, \*\*\**p* = 0.0003; day 1 vs 9 months old, ns.

We characterized the morphology and mechanics of the dye-loaded click alginate cryogel after dye immobilization. Click alginate cryogels exhibit interconnected micron-sized pores that likely permit rapid fluid influx. ^[17a, 34]^ Fluorescence microscopy of dye-conjugated cryogel sections (**Supporting Figure 5**) revealed an interconnected network with micron-sized pores, confirming that dye immobilization does not compromise the porous architecture. Compression testing showed a progressive increase in force with displacement and yielded a compressive stress of 2.11 ± 0.71 MPa at 1.5 mm displacement (engineering strain of 34.7 ± 3.7 % based on initial cryogel heights of 4.36 ± 0.46 mm), at which irreversible cryogel collapse was observed (**Supporting Figure 6A**). Frequency-sweep rheology showed *G*’ remained higher than *G*’’ over the measured frequency range, with *G*’ of approximately 100 Pa and *G*’’ of approximately 10 Pa (**Supporting Figure 6B**), indicating that the cryogel behaves as a viscoelastic solid with a predominantly elastic response. The approximately one-order-of-magnitude difference between *G’* and *G’*’, together with the limited frequency dependence of both moduli, indicates that the covalently crosslinked cryogel retained a stable elastic network over the tested frequency range. The low storage modulus reflects the small-strain response of the hydrated macroporous network, whereas the MPa-level stress at 34.7% strain reflects pore collapse and densification upon compression. These mechanical characteristics are consistent with the formulation- dependent behavior of previously reported tetrazine-norbornene click-alginate hydrogels, in which polymer concentration and the ratio of complementary click-functional groups modulate network mechanics. ^[17a, 20a, 20b, 35]^ Together, these data show that dye immobilization preserved a macroporous network that is soft at small strain and withstands densification at large strain.

### 3.3. *In vitro* evaluation of pH-sensing cryogel activity and stability

After covalently anchoring IRDye 680RD in the click cryogel, we investigated the cryogel’s pH- sensing properties and stability. The dye-conjugated cryogel showed a pH-dependent fluorescence increase in citrate buffer and in protease-containing SGF similar to the dye- tetrazine conjugate (**Figure 5A, 5B**). The similarity of the pH-dependent fluorescence profiles indicates that the conjugation to the alginate scaffold does not distort the dye’s intrinsic pH sensitivity and that the anchored probe remains responsive under gastric fluid-simulating conditions. We next quantified dye release in SGF and SIF (**Figure 5C**). After 24 h, 1.15% and 1.23% of the dye were released in simulated gastric and intestinal fluid, respectively, which indicates a low risk of gastrointestinal absorption. The high stability of dye-conjugated click alginate cryogel reflects the resistance of click bonds and alginate to acidic pH and protease cleavage reported in the literature. ^[17b, 21, 36]^ Finally, cryogels stored for 9 months at 4 °C (in Milli- Q water, protected from light) showed no significant signal loss relative to fresh preparations (**Figure 5D**), indicating that pH-sensing performance is preserved over extended storage in solution.

### 3.4. *In vivo* evaluation of pH-sensing cryogel activity

To evaluate the dye-conjugated cryogel’s capacity to detect gastric pH differences *in vivo*, we first gavaged cryogel-containing buffered solutions at pH 2.0 or pH 5.0 to healthy rats followed by NIR fluorescence imaging. This experiment served as a proof of concept to establish that the cryogel produces a detectable, pH-dependent transabdominal signal *in vivo*; because the cryogel was administered in buffer at the target pH, the pharmacological model below was used to test discrimination of endogenous gastric pH. Consistent with the *in vitro* study, a pH- dependent gastric NIR signal was detected: the gastric fluorescence signal at pH 2.0 was significantly lower than at pH 5.0 (**Figure 6, Supporting Figure 7**), confirming that the dye- conjugated cryogel detects pH differences in the stomach *in vivo* at clinically relevant pH values.

**Figure 6.**
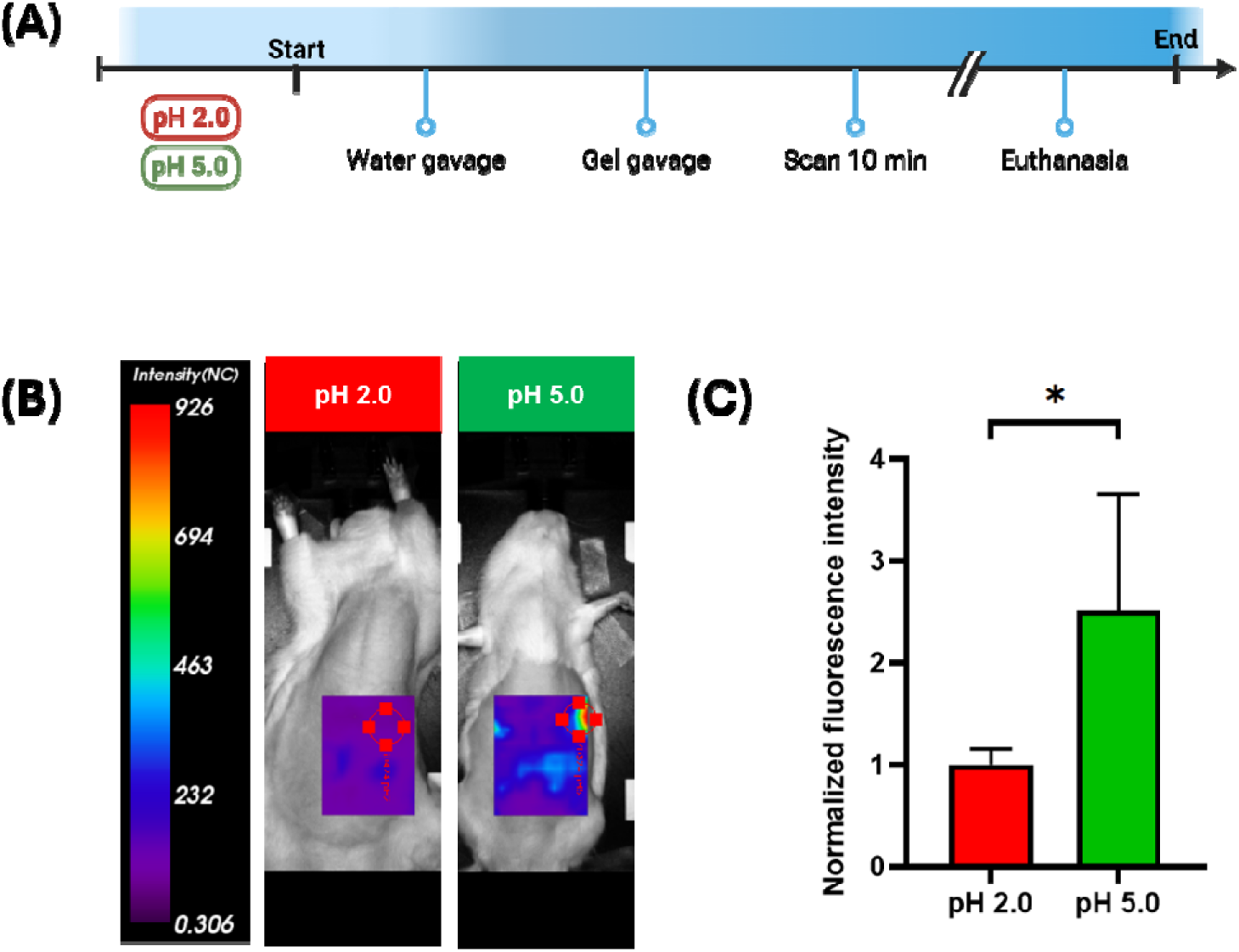
*In vivo* testing of pH-sensitive IRDye 680RD-conjugated click alginate cryogel i healthy rats receiving buffered solutions at pH 2.0 or pH 5.0. (A) Overview of the experimental timeline. (B) Representative image of the fluorescence scans of each group. (C) Gastric fluorescence intensity of rats receiving the cryogel at pH 2.0 and pH 5.0 by gavage. Bar graph results as mean ± SD (*n* = 5). Two-tailed paired *t*-test; \**p* = 0.0388

We next examined whether the dye-conjugated cryogel could distinguish a physiological from a pharmacologically elevated gastric pH in a model of proton-pump inhibitor therapy. Proton-pump inhibition is expected to suppress gastric acid secretion and increase gastric fluid pH in rats. In the literature, oral omeprazole (30 mg/kg/day, 7 days) decreased gastric H^+^ production and increased mean gastric fluid pH from 2.4 in controls to 7.2 in treated rats. ^[37]^ Other studies confirm that omeprazole increases gastric pH in fasted rats and that 20 mg/kg omeprazole potently suppresses gastric acid secretion across oral, intraperitoneal, and intravenous administration routes. ^[38]^ Esomeprazole is the *S*-enantiomer of omeprazole; both enantiomers are converted in the acidic compartment of the parietal cell into the same achiral active inhibitor of the *H^+^, K^+^*-ATPase, ^[39]^ and esomeprazole and omeprazole inhibited gastric acid secretion with similar efficacy in rats *in vivo*. ^[40]^ Doses established for omeprazole in rats were therefore used for esomeprazole. In this study, rats were treated with daily subcutaneous injections of esomeprazole (20 mg/kg; 20 mg/mL in saline; 1 mL/kg) or saline, then imaged after oral administration of the cryogel. Gastric fluorescence in esomeprazole-treated rats was significantly higher than in saline controls (**Figure 7, Supporting Figure 8**). This *in vivo* proof-of-concept study shows that the dye-conjugated click alginate cryogel can discriminate between physiologically acidic and pharmacologically alkalinized gastric environments *in vivo*, supporting its further development toward non-invasive gastric pH monitoring.

**Figure 7.**
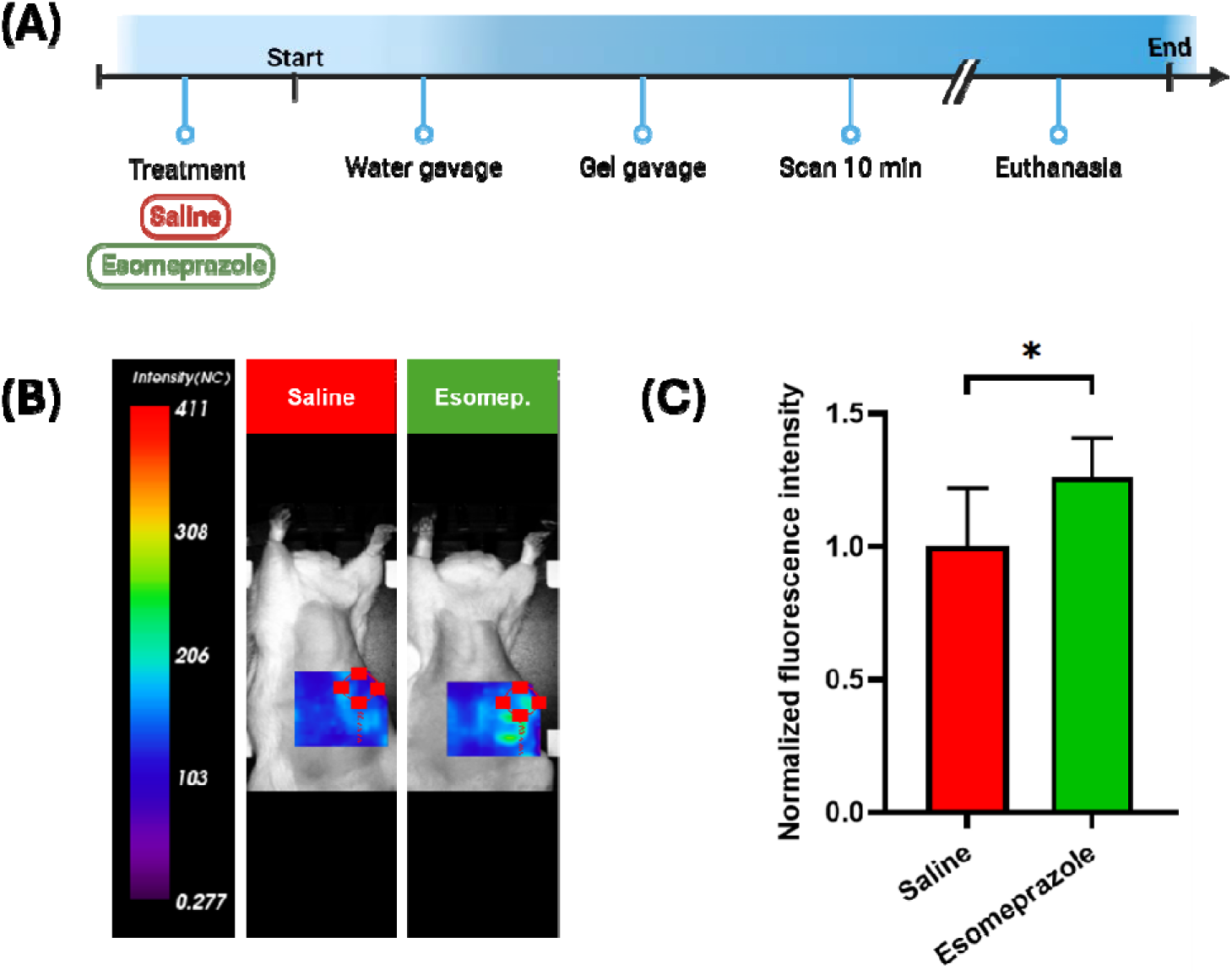
*In vivo* testing of pH-sensitive IRDye 680RD-conjugated click alginate cryogel i rats receiving esomeprazole or saline. (**A**) Overview of the experimental timeline. (**B**) Representative fluorescence images of saline control rats and rats treated with esomeprazole by subcutaneous injection. (**C**) Gastric fluorescence intensity of saline control or esomeprazole- treated rats receiving the cryogel by gavage. Bar graph results as mean ± SD (*n* = 7 per group). Two-tailed unpaired *t*-test; \**p* = 0.0230

### 3.5. *Ex vivo* evaluation of pH-sensing cryogel activity using photoacoustic imaging

To address the relatively low tissue penetration of NIR fluorescence, we further tested the pH- sensitive cryogel using photoacoustic imaging, which combines optical absorption contrast with ultrasound detection and enables imaging across several centimeters of tissue. ^[41]^ Indeed, the IRDye 680RD absorbance at 680 nm makes this dye highly suited for photoacoustic imaging. ^[42]^ The dye-loaded cryogel was gavaged to mice in buffers at pH 2.0 or 5.0, and the stomach was imaged *ex vivo* where significantly different signals for the two pH values were observed (**Supporting Figure 9**), supporting photoacoustic imaging as a complementary readout with potential for deeper-tissue detection.

### 3.6. *Ex vivo* evaluation of cryogel safety profile and dye biodistribution

To assess gastric tissue morphology following repeated oral administration, H&E-stained gastric sections from three saline-treated rats, three esomeprazole-treated rats, and one control rat were evaluated by blinded semi-quantitative histopathology performed by a pathologist (**Supporting Figure 10 and Supporting Table 1**). Gastric mucosal architecture was preserved across all evaluated samples, with intact surface epithelium and no evidence of necrosis **(Supporting Figure 10A-G**). Mild diffuse inflammatory-cell infiltrates, predominantly lymphocytic with occasional eosinophils and neutrophils, were observed in six of the seven animals, including the control, mainly in the deep mucosa and along the gastric glands. One esomeprazole-treated rat exhibited moderate diffuse inflammatory-cell infiltration, predominantly eosinophilic and neutrophilic, extending into the mucosa and submucosa (**Supporting Figure 10F, I, left**). Mild focal parietal-cell alterations, characterized by cellular swelling, granular cytoplasm, and occasional clear cytoplasmic spaces, were observed in one saline-treated rat (**Supporting Figure 10B, H, left**) and were absent from the remaining animals. These findings occurred in individual animals and do not establish treatment-related differences. Overall, these histological findings revealed preserved gastric mucosal architecture without epithelial injury or necrosis, with mild focal parietal-cell alterations in one animal and moderate inflammatory-cell infiltration in another.

To assess potential absorption and extra-gastric biodistribution, excised organs were analyzed by whole-organ fluorescence imaging and quantification (**Supporting Figure 3**). After excision and before imaging, the stomachs were rinsed to remove residual luminal contents. Fluorescence intensities in cryogel-treated animals were comparable to those of the untreated control across the evaluated extra-gastric organs, including the small and large intestines (**Supporting Figure 3K**). The residual fluorescence observed in the rinsed stomach seems higher than in the control. This may be related to the mucoadhesive properties of alginate. ^[17a, 43]^ Plasma fluorescence was comparable among untreated control, esomeprazole-treated, and saline-treated rats (**Supporting Figure 3L**), with no apparent increase following cryogel administration. Together, these findings indicate that oral administration of the cryogel did not seem to lead to an apparent increase in extra-gastric or circulating fluorescence, suggesting limited dye absorption.

## 4. Conclusion and outlook

In summary, we have transformed click alginates from mechanical and delivery scaffolds into a biosensing material for the gastrointestinal tract. A single bioorthogonal chemistry performs both the structural and the functional role: tetrazine-norbornene crosslinking assembles a macroporous cryogel that withstands gastric acid and proteases, while the norbornene groups left in excess covalently anchor IRDye 680RD, the one near-infrared dye in our screen responsive across the clinically decisive pH 2-5 window. This design yields a minimally releasing sensor, with dye release below 1.5% in simulated gastric and intestinal fluids and a pH response retained after nine months of storage. Orally administered cryogels resolved pH 2.0 from pH 5.0 and distinguished esomeprazole- from saline-treated rats by transabdominal near-infrared fluorescence imaging, with extra-gastric organ and plasma fluorescence comparable to untreated controls and preserved gastric mucosal architecture.

The modularity of this architecture is its central strength. Because any tetrazine-bearing reporter couples to the same excess norbornene, the click alginate cryogel becomes a platform technology in which the scaffold is fixed and the analyte is set by the reporter probe. A ROS- sensitive fluorescent dye would report the oxidative mucosal injury that accompanies NSAID gastropathy and chronic gastritis; a crown ether-fluorophore conjugate would bind the ammonium generated by *Helicobacter pylori* urease, giving a non-invasive readout of infection; and aptamers, clicked through the same handle, would extend recognition to ingested toxins such as mycotoxins. The same handle also admits an internal reference: anchoring one of the pH-insensitive near-infrared dyes identified here alongside IRDye 680RD would enable ratiometric, quantitative pH readout. The mucoadhesive character of alginate, alone or combined with gastroretentive dosage forms, would extend the window over which such analytes can be followed from a single administration. Coupled to a portable transabdominal detector, such cryogels could move gastric acid measurement from specialized procedure rooms to the point of care, allowing acid-suppressive therapy to be titrated to the chemistry of the stomach rather than to symptoms. More broadly, covalently functionalized cryogels establish a general strategy for biosensing in physiological environments that dissolve, digest, or leach conventional hydrogels.

## 5. Acknowledgements

Graphical illustrations were created using BioRender, data plots were generated using GraphPad Prism (version 8.0.1), and chemical reaction schemes were drawn using ChemDraw (version 25.5.0). K.C. gratefully acknowledges doctoral scholarships from the Faculté de Pharmacie at Université de Montréal (Bourse du Centenaire and Bourse de la Montagne). This work was supported by a NOVA grant from the Fonds de recherche du Québec - Nature et technologies (FRQNT) and the Natural Sciences and Engineering Research Council of Canada (NSERC) (grant no. 365911, to S.M. and G.Z.), and by a Chercheur-Boursier Junior 1 award from the Fonds de recherche du Québec - Santé (FRQS) (to S.M.). The authors thank Firas El- Mortada (Université de Montréal) for support with hydrogel characterization; Junzheng Peng, M.D., Ph.D. (CrCHUM), for assistance with *in vivo* experimentation; and Mélanie Tremblay, M.Sc. (CrCHUM), for assistance with animal protocols.

## 6. Data availability Statement

The data that support the findings of this study are available from the corresponding author upon reasonable request.

## 8. ToC Figure

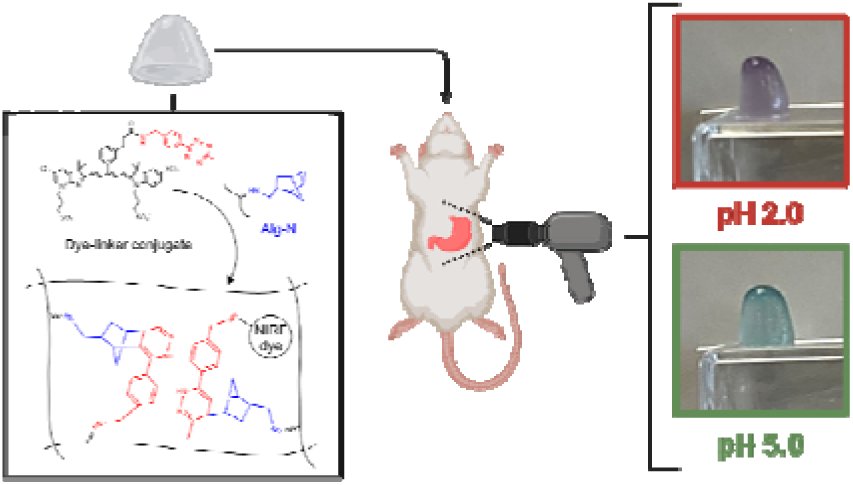

One bioorthogonal chemistry builds the scaffold and the sensor: tetrazine-norbornene crosslinks form an acid- and protease-stable macroporous cryogel, where excess norbornene anchors a near-infrared pH-sensitive dye, enabling rapid sensing and minimal release. Administered orally to rats, the click-alginate cryogel reports gastric pH transabdominally and distinguishes physiological from proton-pump-inhibitor-elevated acidity.

## Supporting information

**Supporting Figure 1.**
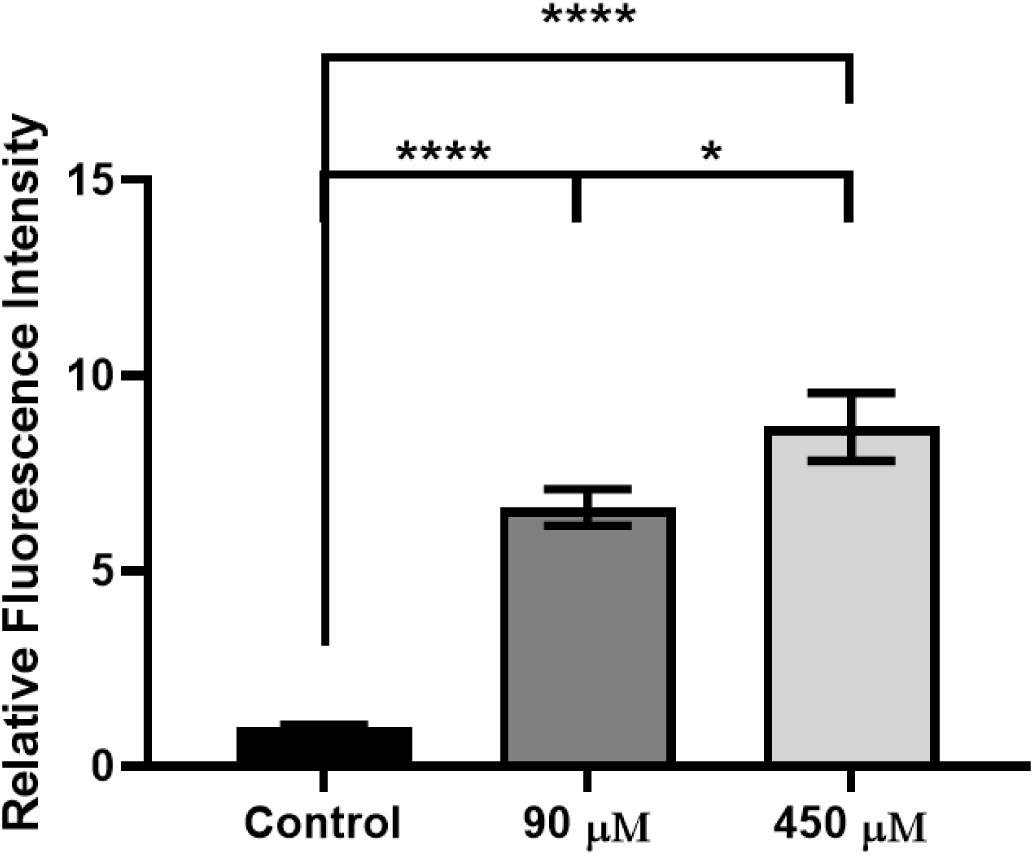
Fluorescence of IRDye 680RD-conjugated cryogel. Relative fluorescence intensity of a native click alginate cryogel (negative control) and cryogels conjugated using 90 µM or 450 µM dye stock solutions. Bar graph results are presented as mean ± SD (*n* = 3). One-way ANOVA with Tukey’s multiple-comparisons test; \*\*\*\**p* < 0.0001, \**p* = 0.0102.

**Supporting Figure 2.**
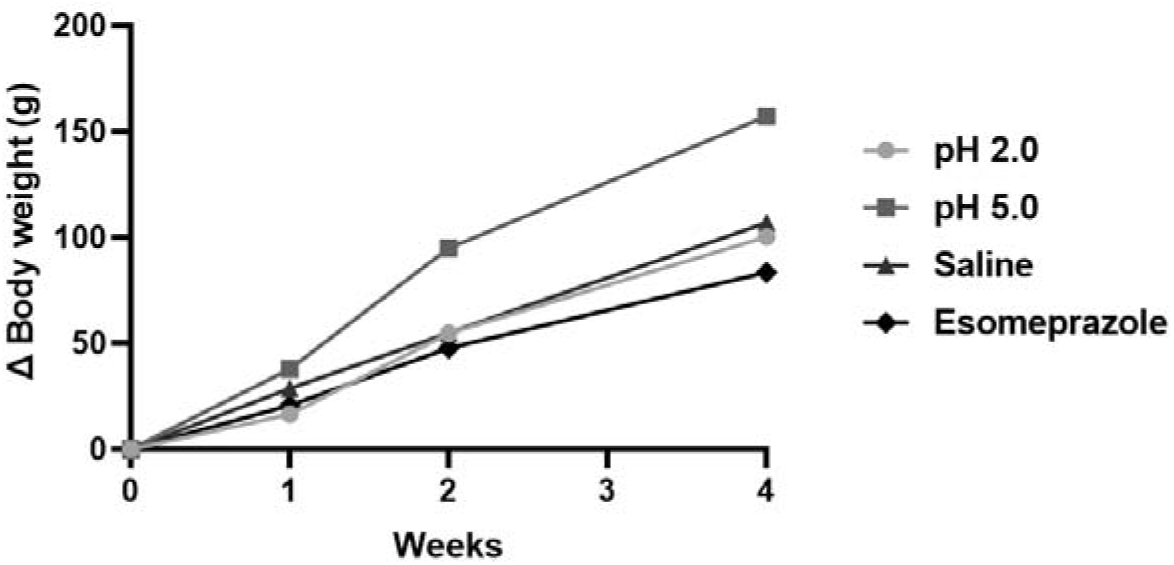
Body weight change during repeated *in vivo* imaging. Change in body weight relative to baseline for individual rats in the pH 2.0 versus pH 5.0 and esomeprazole versus saline studies. Each point represents the body weight of one rat at the indicated timepoint, expressed relative to its baseline body weight. Lines connect repeated measurements from the same animal. No animal showed body-weight loss during the monitoring period.

**Supporting Figure 3.**
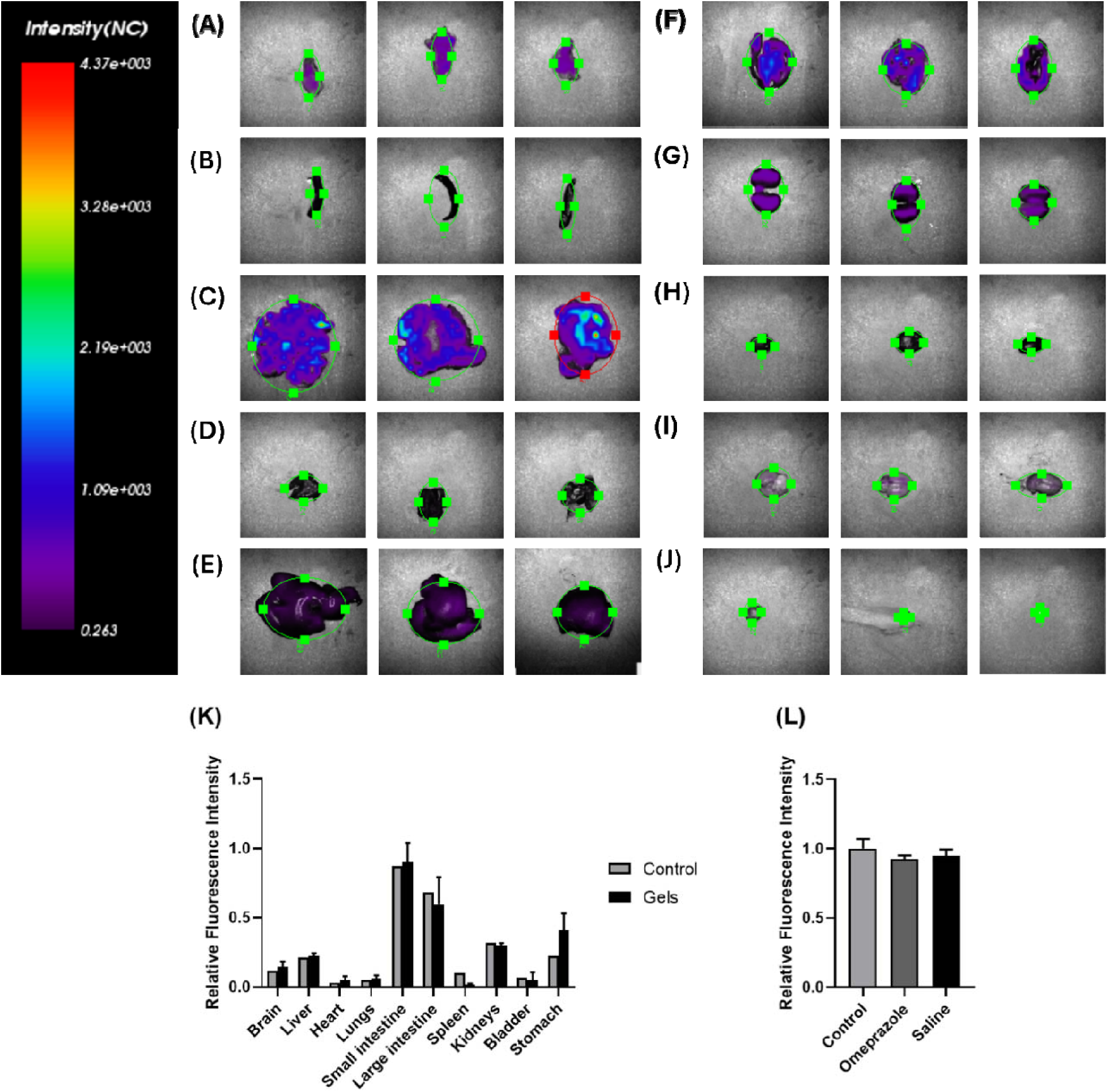
*Ex vivo* fluorescence imaging and quantitative fluorescence analysis after oral administration of dye-conjugated click-alginate cryogels. **(A-J)** Representative ex vivo fluorescence imaging of excised organs from the pH 2.0 versus pH 5.0 study. From left to right, images show one untreated control rat and two cryogel-treated samples. **(A)** Stomach, **(B)** spleen, **(C)** small intestine, **(D)** lungs, **(E)** liver, **(F)** large intestine, **(G)** kidneys, **(H)** heart, **(I)** brain, and **(J)** bladder. **(K)** Relative fluorescence intensity of excised organs from the pH 2.0 versus pH 5.0 study in rats. Grey bars represent an untreated, age- matched healthy control rat (*n* = 1), and black bars represent cryogel-treated animals (*n* = 2). **(L)** Relative plasma fluorescence intensity in an untreated, age-matched healthy control rat that never received a cryogel, and in cryogel-gavaged esomeprazole-treated and saline-treated rats *(n* = 5). All data are presented as mean ± SD. No statistical testing was performed (descriptive comparison).

**Supporting Figure 4.**
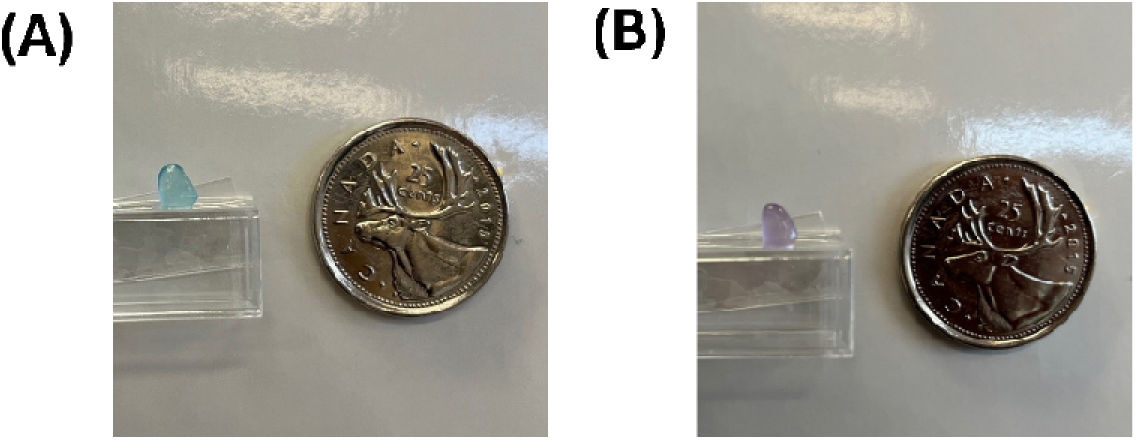
Macroscopic pH-responsive color change of IRDye 680RD- conjugated click alginate cryogels. Photographs of cryogels loaded with IRDye 680RD. Fluorescence imaging formulation equilibrated at pH 5.0 **(A)** or pH 2.0 **(B);** a Canadian 25 cent coin is included for scale.

**Supporting Figure 5.**
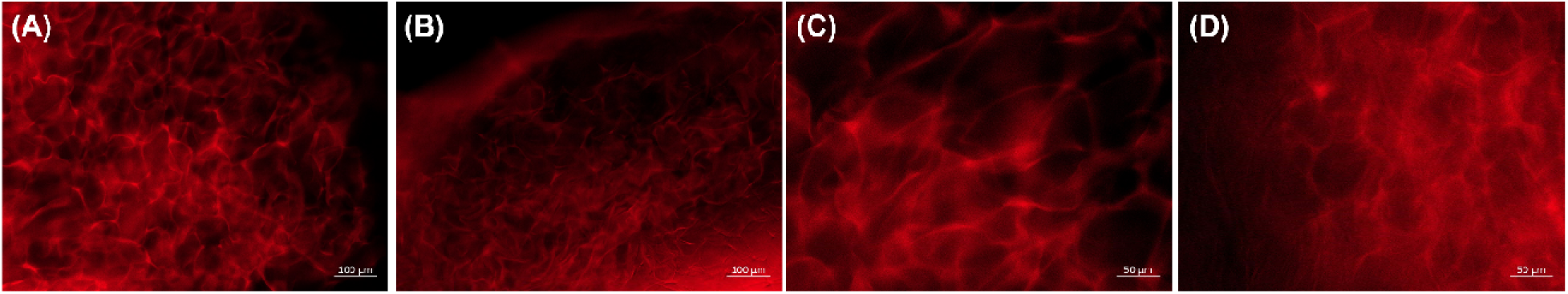
Macroporous structure of the dye-conjugated click alginate cryogel. Fluorescence images of cryogel sections showing the macroporous architecture at (**A, B)** 10 magnification (100 µm scale bars) and **(C, D)** 20 × magnification (50 µm scale bars).

**Supporting Figure 6.**
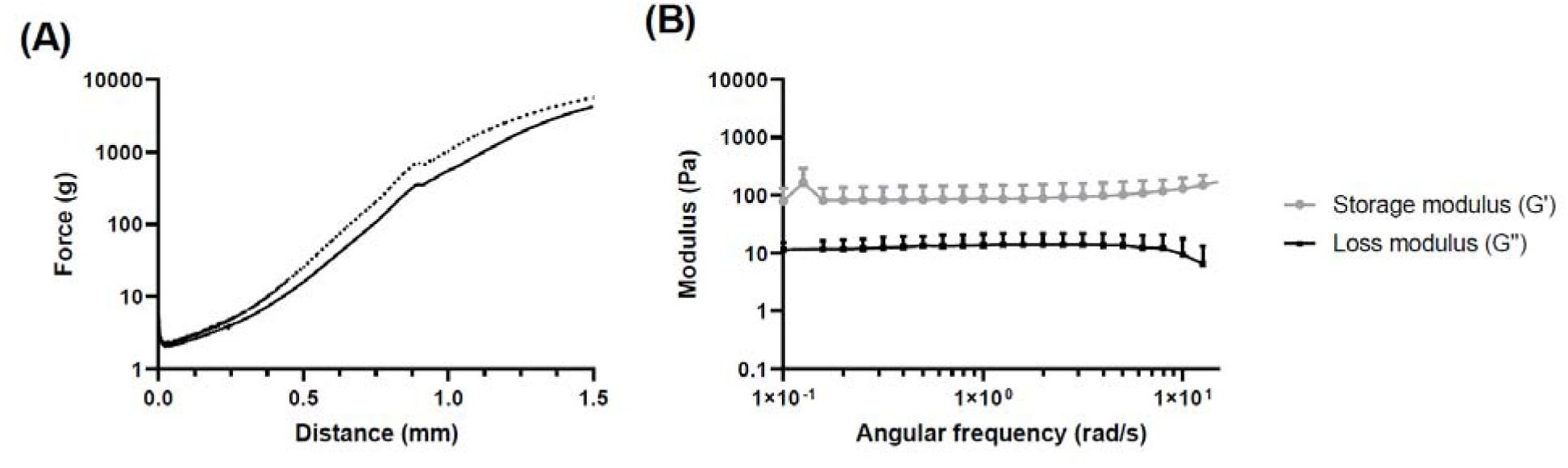
Mechanical and rheological characterization of the dye-conjugate click alginate cryogel. **(A)** Force-distance compression profile (solid line, mean; dotted line, SD). **(B)** Storage (*G*′) and loss (*G*″) moduli as a function of angular frequency (ω = 2πf; 0.1-10 rad/s, ≈ 0.016-1.6 Hz). Data are mean ± SD (*n* = 3).

**Supporting Figure 7.**
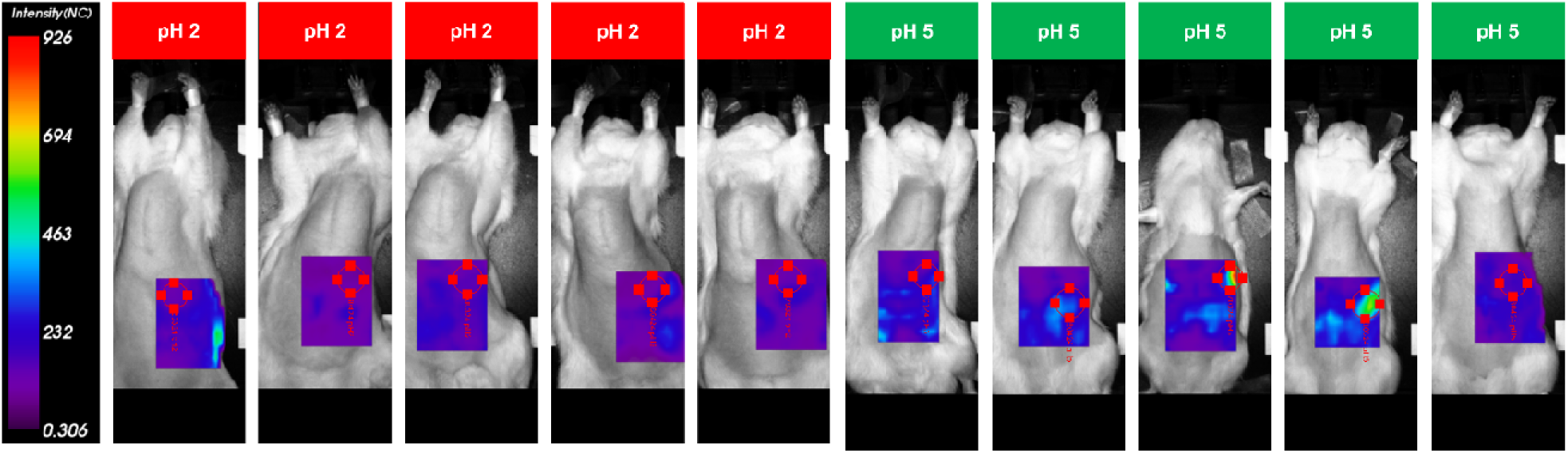
*In vivo* testing of pH-sensitive IRDye 680RD-conjugated click alginate cryogel in healthy rats receiving buffered solutions at pH 2.0 or pH 5.0. Compilation of fluorescence imaging scans of all images in Figure 6.

**Supporting Figure 8.**
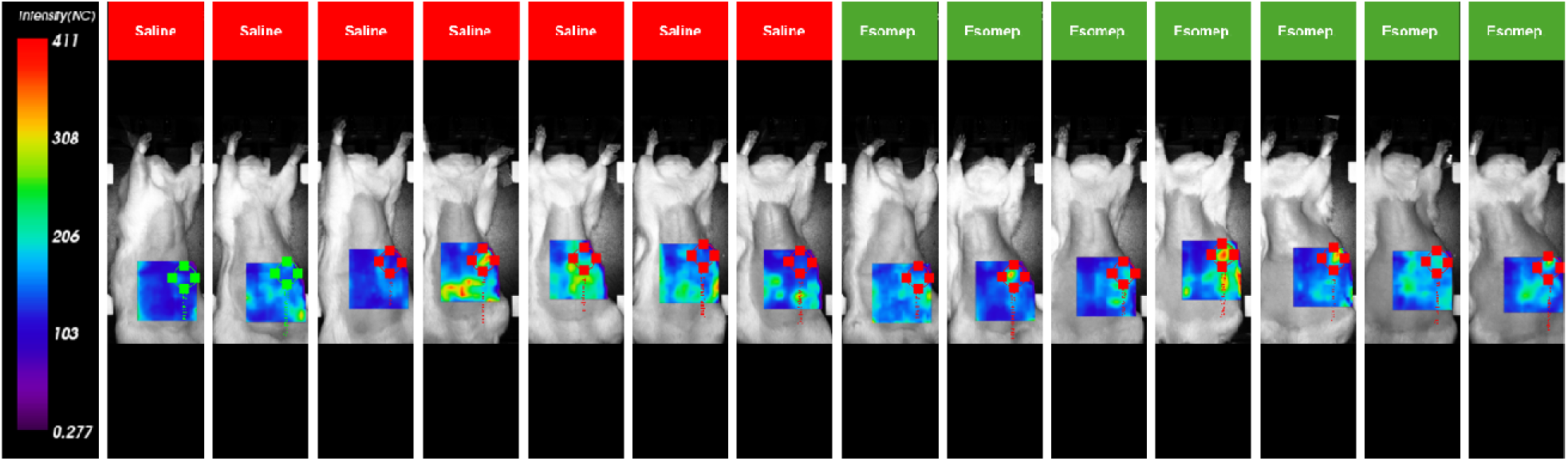
*In vivo* testing of pH-sensitive IRDye 680RD-conjugated click alginate cryogel in rats receiving esomeprazole or saline. Compilation of fluorescence imaging scans of all images in Figure 7.

**Supporting Figure 9.**
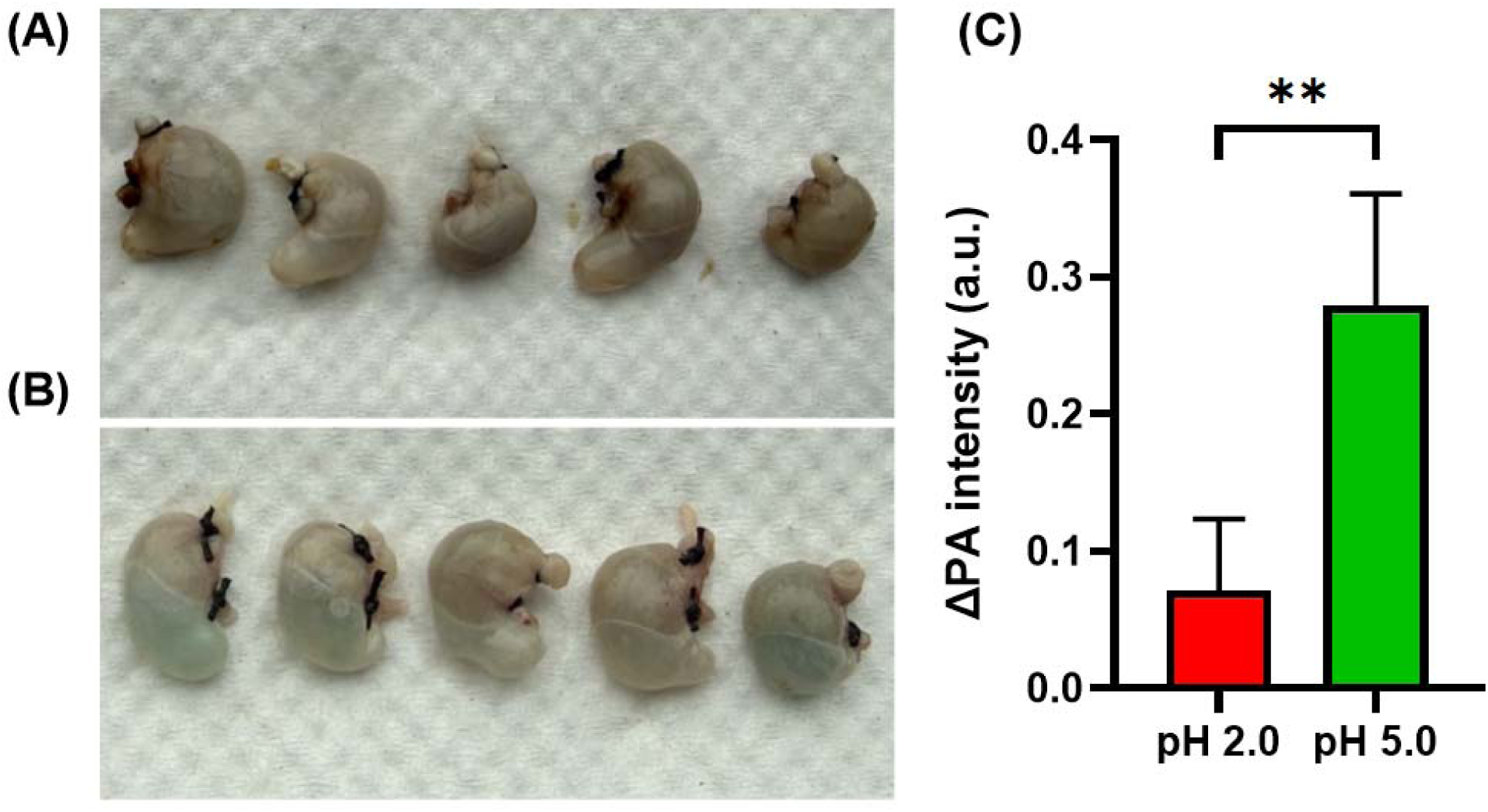
*Ex vivo* testing of pH-sensitive IRDye 680RD-conjugated click alginate cryogel in healthy mice receiving buffered solutions at pH 2.0 or pH 5.0 usin photoacoustic imaging. Representative photographs of excised mouse stomachs following administration of dye-loaded click alginate cryogels at **(A)** pH 2.0 or **(B)** pH 5.0. **(C)** Average photoacoustic (PA) signal measured from gastric regions of interest following excitation at 680 nm, after subtraction of the PA signal of buffer-only negative controls at the corresponding pH. Data are presented as mean ± SD (*n* = 5 mice per group). Two-tailed unpaired *t*-test; \**p* = 0.0116

**Supporting Figure 10.**
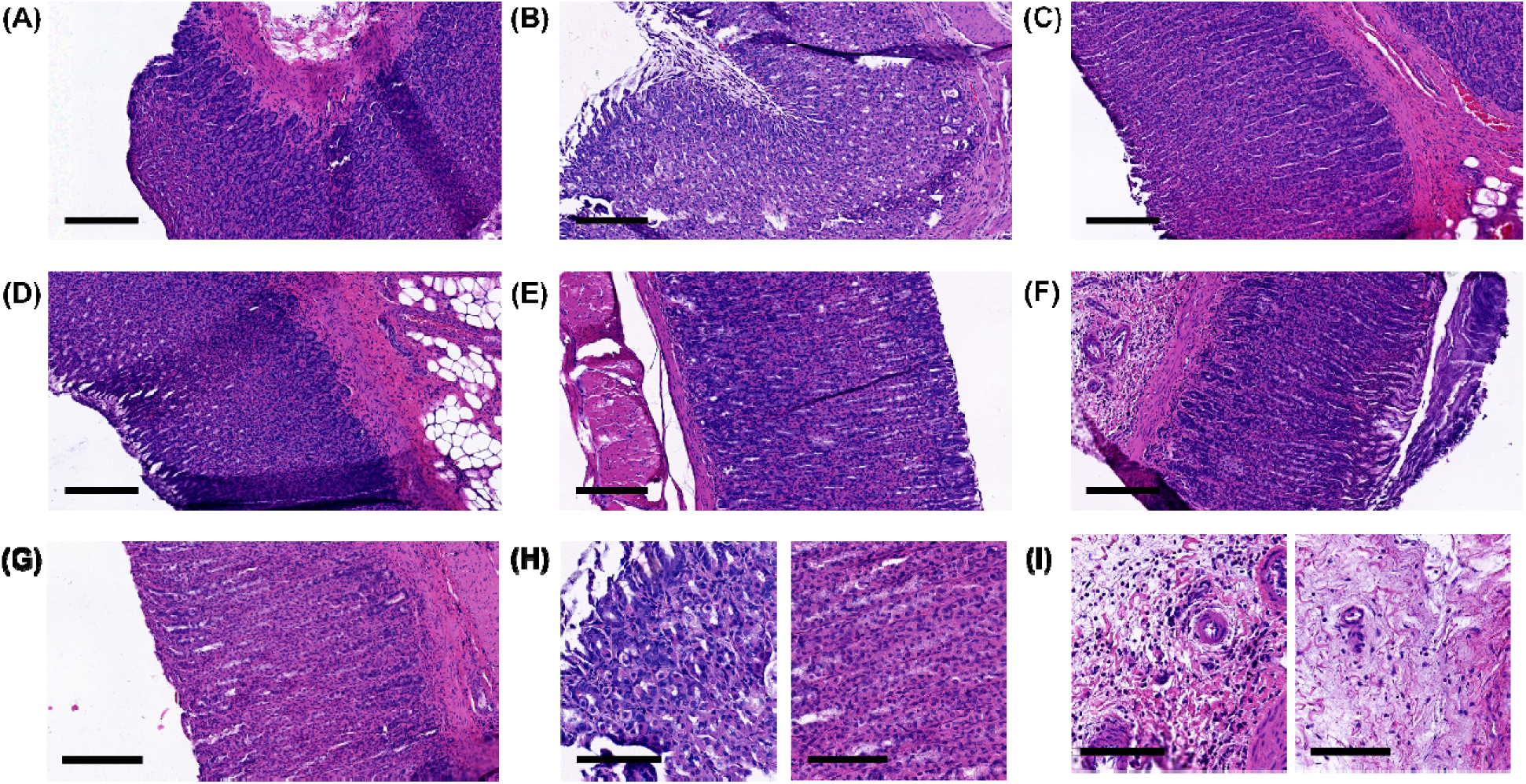
Gastric histology following oral gavage in saline-treated, esomeprazole-treated, and control rats. **(A-C)** Representative H&E-stained gastric sections from three saline-treated rats**. (D-F)** Representative H&E-stained gastric sections from three esomeprazole-treated rats**. (G)** Representative H&E-stained gastric section from the control rat**. (H)** Higher-magnification images of the same stomachs shown in (B) (left, saline-treated) and (G) (right, control), illustrating mild focal parietal-cell alterations observed in one saline-treated rat and their absence in the control specimen. **(I)** Higher-magnification images of the same stomachs shown in (F) (left, esomeprazole-treated) and (C) (right, saline-treated), illustrating moderate inflammatory- cell infiltration observed in one esomeprazole-treated rat and mild infiltration in the saline-treated specimen. Scale bars: (A-G) 400 µm; (H, I) 200 µm.

**Supporting Table 1.** Gastric histopathology scores. H&E-stained gastric sections were anonymized before blinded semi-quantitative evaluation. Scores range from 0 to 3: 0 = absent/no abnormality; 1 = mild/focal alteration; 2 = moderate/multifocal or diffuse alteration; 3 = severe/diffuse alteration. Epithelial integrity reflects preservation of gastric epithelium and the presence and extent of superficial erosions. Individual animal scores are reported.

| **Group** | **Epithelial integrity** | **Parietal-cell alteration** | **Inflammation** | **Necrosis** |
|---|---|---|---|---|
| Control, untreated | 0 | 0 | 1 | 0 |
| Saline | 0 | 0 | 1 | 0 |
|  | 0 | 1 | 1 | 0 |
|  | 0 | 0 | 1 | 0 |
| Esomeprazole | 0 | 0 | 1 | 0 |
|  | 0 | 0 | 1 | 0 |
|  | 0 | 0 | 2 | 0 |

